# An Infected Cryopreserved Precision-Cut Lung Slice Model for Drug Susceptibility Testing in *Pneumocystis murina* Pneumonia

**DOI:** 10.64898/2026.11.19.767844

**Authors:** Madeline Pellegrino, Jenna L. McLean, Theodore J. Kottom, Andrew H. Limper

## Abstract

Precision-cut lung slices (PCLS) preserve native three-dimensional lung architecture and cellular complexity, providing a valuable ex vivo platform for studying pulmonary host-pathogen interactions. However, existing PCLS infection models typically introduce microorganisms into previously uninfected tissue, while fresh PCLS prepared from infected lungs have limited experimental flexibility and scalability. To our knowledge, cryopreserved PCLS generated from lungs with an established in vivo pulmonary infection have not been described. Herein, we evaluated cryopreserved PCLS from mice with established *Pneumocystis murina* pneumonia (PCP) as an ex vivo platform for infection and therapeutic studies. PCLS were prepared from dexamethasone-immunosuppressed mice with established PCP, rested for 24 h, cryopreserved in a novel freezing medium for 12 days in liquid nitrogen, rapidly thawed, recovered for 24 h, and treated with pentamidine for 72 h. Viability was assessed in uninfected and PCP-infected PCLS before and after cryopreservation. Although post-thaw viability decreased in both groups, substantial viability was still retained, allowing further experimentation. qPCR showed that cryopreservation did not alter expression of the type I and type II alveolar epithelial markers podoplanin (*Pdpn*) and surfactant protein C (*Sftpc*) in either group. Importantly, pentamidine significantly reduced *Pneumocystis* mitochondrial small subunit (*mtSSU*) rRNA levels in thawed PCP-infected PCLS, demonstrating suitability for anti-*Pneumocystis* drug testing. These findings establish the first cryopreserved ex vivo PCP model using infected mouse PCLS, preserving endogenous fungal burden and infection-associated host responses while enabling flexible, scalable studies of *Pneumocystis* biology, host-pathogen interactions, therapeutic efficacy, and potentially other pulmonary infections.

## INTRODUCTION

Pneumocystis pneumonia (PCP) in humans is caused by *Pneumocystis jirovecii* (1), a member of the phylum Ascomycota and a fungus closely related to *Saccharomyces cerevisiae* and *Schizosaccharomyces pombe* (2). Different *Pneumocystis* species infect different mammals (3, 4), including *P. murina* in mice and *P. carinii* in rats (5). Ubiquitous in nature, PJP occurs in patients with T-cell deficiencies, including those with HIV infection, cancer, and solid organ transplants (6–8). *Pneumocystis jirovecii* pneumonia (PJP) continues to impose a significant global health burden (7), therefore, new methods to study *Pneumocystis* biology and develop therapeutics are needed. Progress in developing novel therapeutic candidates and understanding the organism’s life cycle has been hindered by the lack of a physiologically relevant system that can recapitulate the lung microenvironment. In vitro culture of *Pneumocystis* is limited by the reliability of these systems and by the organism’s poor viability over extended periods in vitro (9, 10). Animal models that mimic the immunocompromised state are available, but they are costly and only provide low throughput for drug development studies (10).

We and others have recently shown that precision-cut lung slices (PCLS) are an effective, small format ex vivo system for studying *Pneumocystis* permitting higher-throughput studies (11, 12). Unlike conventional in vitro cell-based and cell-free culture systems, PCLS preserve the three-dimensional native architecture of the lung, including intact airways, alveolar spaces, and extracellular matrix (13, 14). In addition, PCLS maintain multiple host myeloid and non-myeloid cell populations (15–18) and, in PCP-derived PCLS, preserve the authentic *Pneumocystis*-infected lung microenvironment (11, 12).

In this report, we expand on our previous work using PCP-infected PCLS (12) and describe a novel PCLS cryopreservation method that maintains lung tissue and *Pneumocystis* fungal viability for at least 12 days after thawing following cryopreservation. Using the anti-*Pneumocystis* drug pentamidine (19), we demonstrate as proof of concept that this cryopreservation model can be used for PCLS downstream future high-throughput therapeutic drug testing, and studies of *Pneumocystis*-host cell interactions. Together, our results advance prior *Pneumocystis* PCLS work from our laboratory and others and provide initial results for a novel cryopreservation ex vivo platform for *Pneumocystis* research.

## METHODS

### Murine Model of *Pneumocystis murina* Infection

Six-week-old male BALB/c mice (Charles River Laboratories) were immunosuppressed by administering drinking water containing antibiotics and 4 mg/mL dexamethasone sodium phosphate beginning one day before infection (20). Frozen *P. murina* inoculum stocks were generated from the lungs of heavily infected donor mice, purified by sequential differential centrifugation and filtration, and quantified to standardize the infectious dose as previously described (21). On the day of infection, mice were anesthetized with isoflurane and inoculated intranasally with 1 x 10 *P. murina* organisms suspended in sterile phosphate-buffered saline (PBS). Infection was allowed to progress for 8-12 weeks while animals remained on dexamethasone- and antibiotic-supplemented drinking water ad libitum to maintain immunosuppression and reduce the risk of secondary bacterial infections (22).

### Cryopreservation and Recovery of Precision-Cut Lung Slices

After the 24 h rest period, both uninfected and PCP infected PCLS were cryopreserved for future use. Briefly, 10 individual slices were transferred to cryovials containing 900 µL of freezing media (DMEM/F12) (1:1) supplemented with 300mM Trehalose and 1X penicillin-streptomycin-amphotericin (23). Samples were then incubated at 37°C, 5% CO_2_ for a minimum of 30 minutes before addition of 100 µL of DMSO to achieve a final concentration of 10%. Cryovials were subsequently moved to a controlled-rate freezing container (Mr. Frosty™, Sigma Aldrich, Inc.) and stored overnight at -80°C before transfer to liquid nitrogen for long-term storage. For thawing, a 6-well plate was prepared with 1 mL of warmed culture media supplemented with 2X antibiotic-antimycotic before cryovials were thawed rapidly in a 37°C water path. The contents of each were poured into individual wells, rinsed with 1 mL, and this process was repeated 4 times. The media in each well was then replaced entirely and the slices transferred to individual wells and allowed to incubate for 24 h at 37°C, 5% CO_2_.

### Assessment of PCLS Metabolic Activity

The metabolic activity of *Pneumocystis*-infected precision-cut lung slices (PCLS) was evaluated using the PrestoBlue™ Cell Viability Reagent (Invitrogen, A13261), a resazurin-based fluorometric assay that measures cellular reducing capacity (11, 24). PrestoBlue™ was diluted to a final concentration of 10% in PCLS culture medium, and 150 µL of the reagent mixture was added to each well of a 96-well plate containing individual lung slices. Wells containing culture medium alone served as background controls. Following a 2 h incubation at 37°C, 100 µL of the reaction medium from each well was transferred to a black-walled 96-well fluorescence plate. Fluorescence intensity was measured using a SpectraMax M2e microplate reader (Molecular Devices) with excitation and emission wavelengths set to 540 nm and 590 nm, respectively. Background fluorescence was corrected by subtracting values obtained from media-only controls using SoftMax^®^ Pro software (version 7.3), and the resulting fluorescence values were used as an indicator of relative tissue metabolic activity, and hence viability.

### RNA Isolation and Quantitative Gene Expression Analysis

Following the completion of culture, *Pneumocystis*-infected precision-cut lung slices (PCLS) were gently blotted to remove residual culture medium and immediately lysed in Buffer RLT Plus (RNeasy^®^ Plus Mini Kit, Qiagen). Tissue disruption was performed using a TissueLyser II (Qiagen) with stainless-steel beads under cold conditions (4°C). Genomic DNA was removed by passing the lysates through gDNA Eliminator spin columns, after which the flow-through was combined with 70% ethanol and loaded onto RNeasy spin columns for RNA purification. Total RNA was isolated according to the manufacturer’s protocol and subsequently further purified and concentrated using an RNA Clean & Concentrator Kit (R1016, Zymo Research).

Complementary DNA (cDNA) was synthesized from purified RNA using the iScript™ Select cDNA Synthesis Kit (Bio-Rad) with the supplied oligo(dT)/random hexamer primer mixture. Quantitative real-time PCR (qRT-PCR) was then carried out on a CFX96™ Real-Time PCR Detection System (Bio-Rad) using SYBR™ Green master mix (Bio-Rad). Transcript levels of *Pdpn*, a marker of alveolar type I epithelial cells, and *Sftpc*, a marker of alveolar type II epithelial cells, were quantified and normalized to the reference gene *B2m* (β2-microglobulin) (11, 12, 25). To assess the *P. murina* burden within infected PCLS after pentamidine treatment (26, 27), fungal mitochondrial small-subunit ribosomal RNA (*mtSSU* rRNA) was quantified by qRT-PCR using an approach adapted from previously published methods (12). Primer sequences for all targets are listed in Supplementary Table 1.

### Statistical Methods

All statistical analyses were performed using GraphPad Prism (version 11.1.0; GraphPad Software, San Diego, CA). Differences among experimental groups were assessed by one-way analysis of variance (ANOVA), with two-tailed Student’s *t*-tests used for pairwise comparisons when appropriate. Data were generated from experiments using lung tissue obtained from three mice, with each treatment group consisting of 3-17 precision-cut lung slices (PCLS), depending on the experimental design. In all figures, each individual data point (black circle) represents a single PCLS. A *P* value of less than 0.05 was considered a statistically significant difference.

## RESULTS

### Cryopreserved Mouse PCLS Retain Sufficient Metabolic Activity for Downstream PCP Studies

Resazurin reduction assays demonstrated a significant decrease in metabolic activity following 12 days of cryopreservation and subsequent thawing in both PCP-infected and uninfected PCLS compared with freshly prepared slices (Fig. 1A). As we previously reported (12), PCP-infected PCLS exhibited significantly greater resazurin reduction than uninfected PCLS before cryopreservation. Despite a significant reduction in metabolic activity following cryopreservation, thawed PCLS retained measurable and reasonable resazurin levels, indicating the presence of viable and metabolically active PCLS samples. Furthermore, treatment of post-thaw PCLS with 1.0 µg/mL pentamidine did not significantly alter resazurin levels in either uninfected or PCP-infected PCLS compared with their respective DMSO-treated controls (Fig. 1B). Together, these results demonstrate that cryopreserved PCLS retain metabolic activity following thawing and remain suitable for downstream ex vivo studies of *P. murina* infection.

**Figure 1.**
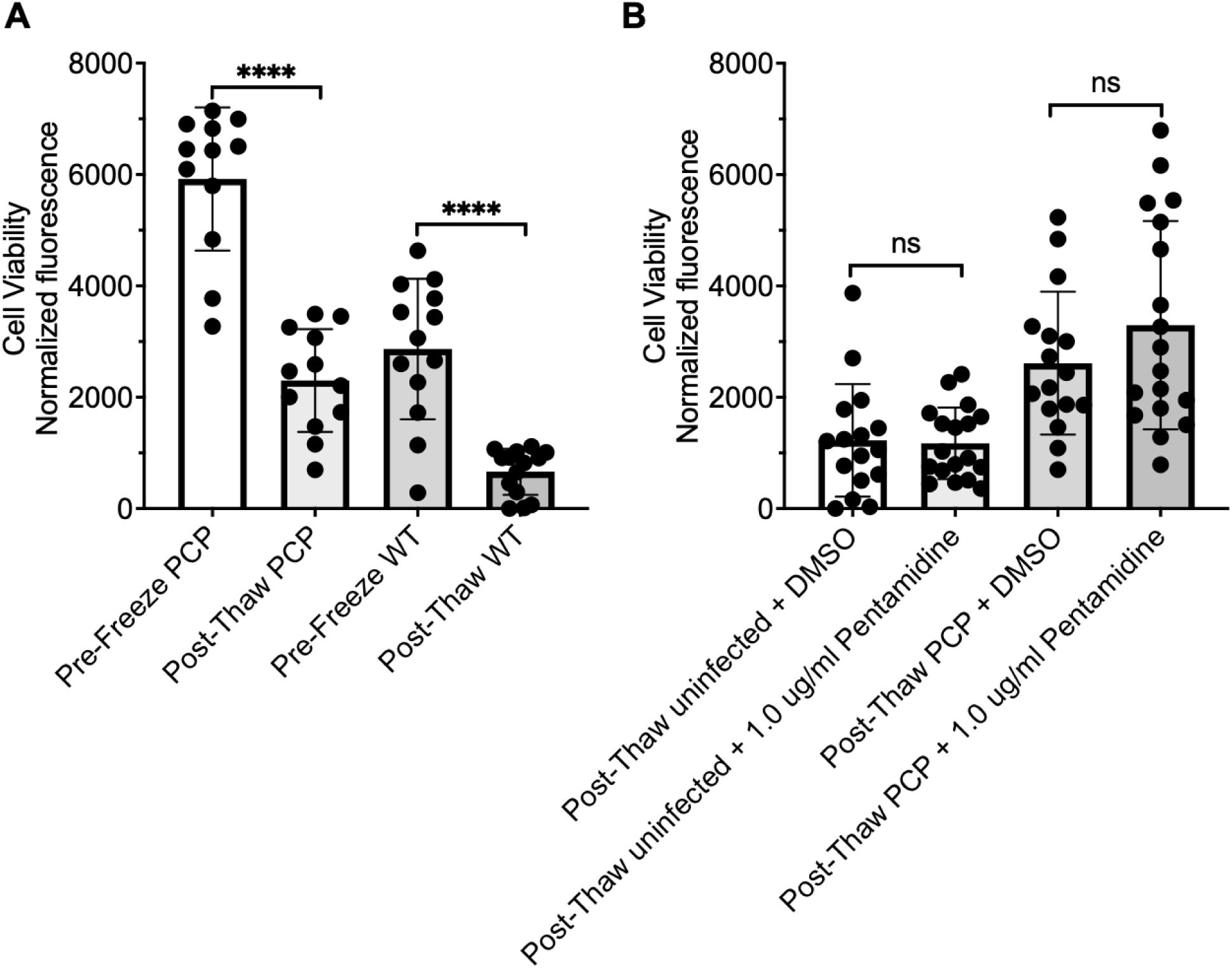
Cryopreservation reduces PCLS metabolic activity but does not alter the metabolic activity of post-thaw PCLS following pentamidine treatment. **(A)** Metabolic activity of freshly prepared (pre-freeze) and cryopreserved (post-thaw) *P. murina*-infected (PCP) and uninfected (WT) precision-cut lung slices (PCLS) was assessed using the resazurin reduction assay. PCLS were cryopreserved for 12 days, thawed, and cultured for 24 h before analysis. Resazurin fluorescence was significantly reduced in post-thaw PCLS compared with freshly prepared PCLS in both infected and uninfected groups. **(B)** Effect of pentamidine treatment on the metabolic activity of post-thaw PCLS. Following thawing and a 24-h recovery period, uninfected and PCP-infected PCLS were cultured for an additional 72 h in the presence of 1.0 µg/mL pentamidine or DMSO vehicle control before assessment by the resazurin reduction assay. Pentamidine treatment did not significantly alter resazurin fluorescence in either uninfected or PCP-infected PCLS. Results are presented as the mean ± SEM. **** *P*< 0.0001; ns, not significant.

### Cryopreservation Does Not Alter Type I or Type II Epithelial Marker Expression in PCLS

Similar to what we have published before in the infected PCP PLCS model (12), *Pdpn* mRNA expression is significantly upregulated in the fungal infected PCLS versus the uninfected PCLS in both pre and post cryopreservation and pentamidine treated PCLS (Figure 2A). *Sftpc* mRNA expression was significantly reduced in PCP-infected PCLS compared with uninfected controls under three of the four experimental conditions (Figure 2B). In the remaining comparison (post-thaw uninfected versus post-thaw PCP-infected PCLS), *Sftpc* expression was also reduced following PCP infection, although this difference did not reach statistical significance (*P* = 0.19).

**Figure 2.**
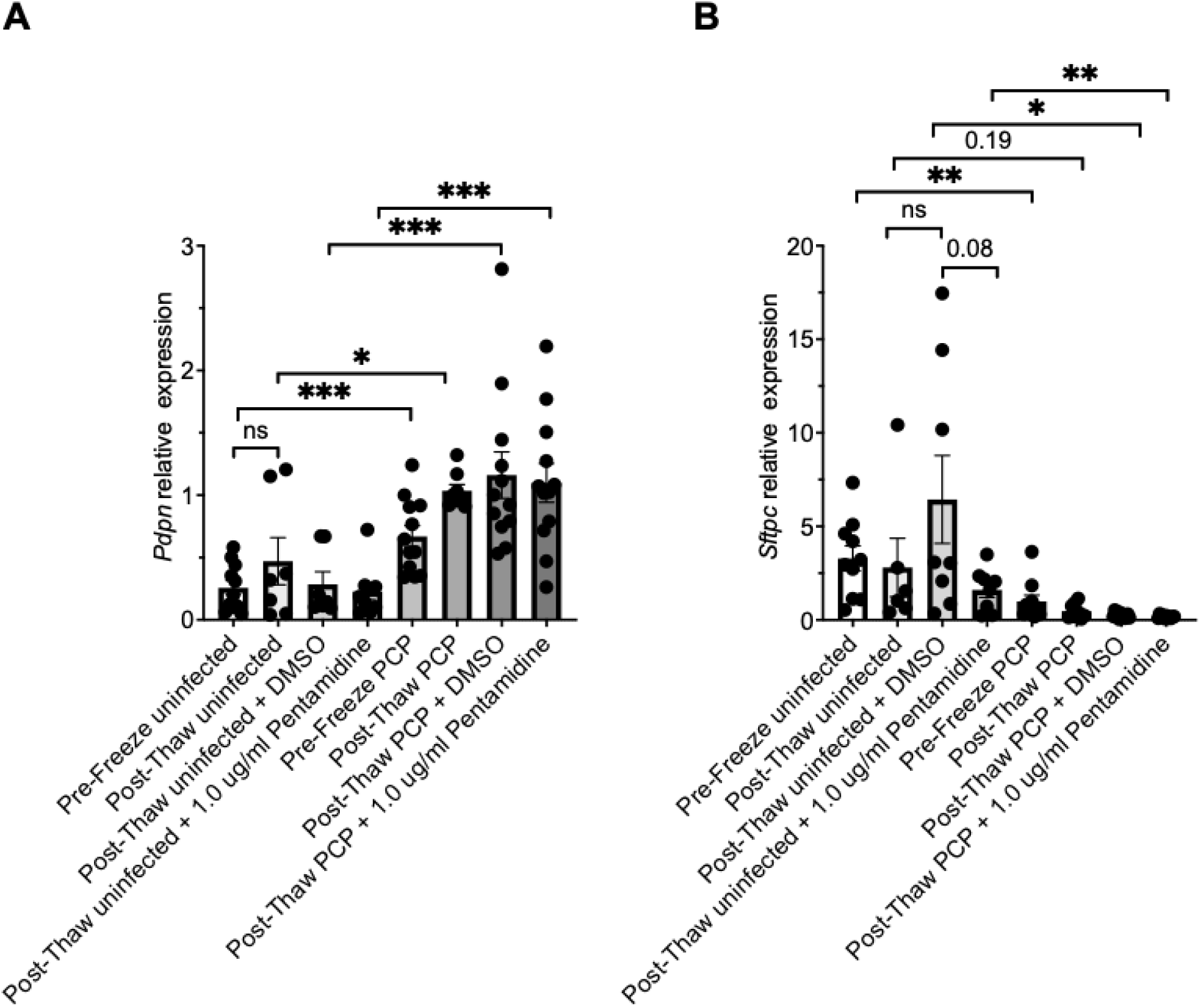
Cryopreservation preserves the expression of *Pdpn* and *Sftpc* mRNA in uninfected PCP-infected PCLS. **(A)** Relative *Pdpn* mRNA expression was quantified by RT-qPCR in freshly prepared (pre-freeze) and cryopreserved (post-thaw) uninfected and *P. murina*-infected (PCP) PCLS. Following thawing and a 24-h recovery period, post-thaw uninfected and PCP-infected PCLS were cultured for an additional 72 h in the presence of 1.0 µg/mL pentamidine or DMSO vehicle control before RNA isolation. *Pdpn* mRNA expression was maintained following cryopreservation and was significantly increased in PCP-infected PCLS compared with uninfected controls. Pentamidine treatment did not significantly alter *Pdpn* expression in either uninfected or PCP-infected PCLS. **(B)** Relative *Sftpc* mRNA expression was quantified under the same experimental conditions. *Sftpc* mRNA expression was significantly reduced in PCP-infected PCLS compared with uninfected controls under three of the four experimental conditions. The remaining comparison (post-thaw uninfected versus post-thaw PCP-infected PCLS) also showed reduced *Sftpc* expression following PCP infection, although this difference did not reach statistical significance. Pentamidine treatment did not significantly affect *Sftpc* mRNA expression in either uninfected or PCP-infected PCLS. Results are presented as the mean ± SEM. *P* values are indicated in the figure; ns, not significant.

### Cryopreserved PCP-infected PCLS Retain Metabolic Activity and Support Antifungal Drug Evaluation

Resazurin reduction assays demonstrated that cryopreservation for 12 days followed by thawing significantly reduced the metabolic activity of PCP-infected PCLS compared with freshly prepared PCP-infected PCLS (Fig. 3A). Despite this reduction, thawed PCLS retained measurable resazurin levels, indicating that the lung slices remained viable and metabolically active following cryopreservation. To validate cryopreserved PCLS as an ex vivo platform for antifungal drug evaluation, post-thaw PCP-infected PCLS were treated with 1.0 µg/mL pentamidine or DMSO (vehicle) for 72 h. Quantification of *P. murina* viability by qPCR targeting the mitochondrial small subunit (*mtSSU*) rRNA demonstrated a significant reduction in viable organisms following pentamidine treatment compared with DMSO-treated controls (Fig. 3B), providing proof-of-concept that cryopreserved PCP-infected PCLS can be used to evaluate antifungal efficacy.

**Figure 3.**
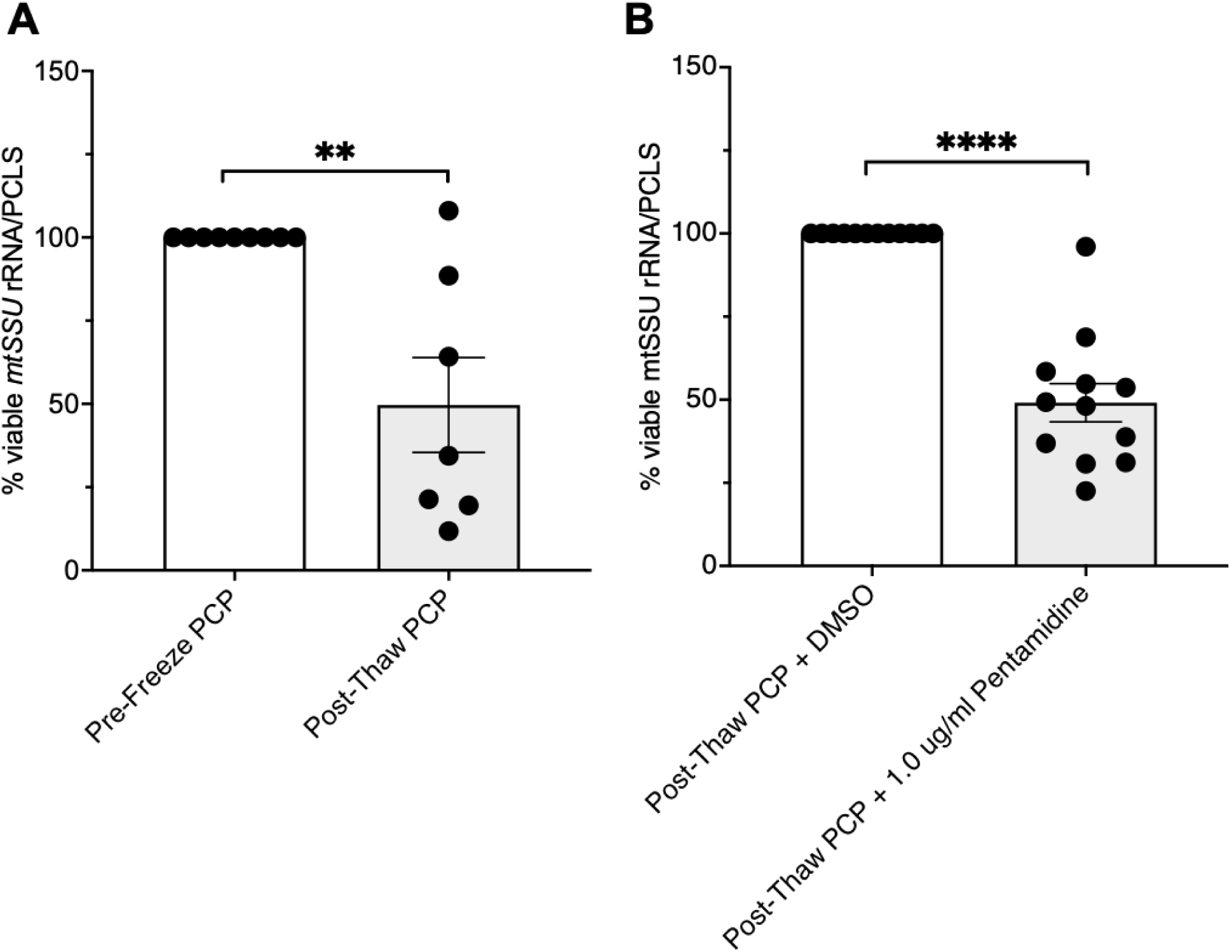
Cryopreserved PCP-infected PCLS retain *P. murina* viability and support antifungal efficacy testing. **(A)** *P. murina* viability was quantified by qPCR targeting mitochondrial small subunit (*mtSSU*) rRNA in freshly prepared (pre-freeze) and cryopreserved (post-thaw) PCP-infected precision-cut lung slices (PCLS). PCLS were cryopreserved for 12 days, thawed, and cultured for 24 h before analysis. Cryopreserved PCP-infected PCLS exhibited a significant reduction in the percentage of viable *P. murina* compared with freshly prepared PCP-infected PCLS. **(B)** To evaluate the suitability of cryopreserved PCLS for antifungal efficacy studies, post-thaw PCP-infected PCLS were cultured for 24 h in media and then an additional 72 h in the presence of 1.0 µg/mL pentamidine or DMSO vehicle control. *P. murina* viability was quantified by qPCR targeting *mtSSU* rRNA. Pentamidine treatment significantly reduced the percentage of viable *P. murina* compared with DMSO-treated controls. Results are presented as the mean ± SEM. *P* < 0.01, **** *P* < 0.0001.

## DISCUSSION

Precision-cut lung slices (PCLS) provide an attractive ex vivo platform for investigating host-pathogen interactions because they preserve the native three-dimensional architecture of the lung and maintain the complex cellular composition that is lost in conventional monolayer cultures and absent from many in vitro model systems. We and others have previously used this platform to investigate host-lung interactions during *Pneumocystis* infection, while others have employed PCLS to study host responses to bacterial and viral pathogens (11, 12, 28, 29).

Previous studies have also demonstrated that cryopreserved PCLS remain viable and functional for applications including airway physiology (30), toxicology (31), immune response studies (32), and pharmacological testing (33). For example, Rosner et al. showed that cryopreserved murine PCLS maintained airway contractile responses following thawing (34), while Patel et al. developed an on-demand cryopreserved human PCLS platform that maintained tissue viability and immune competence for toxicological applications (35). However, these studies focused primarily on host tissue function and did not evaluate cryopreserved PCLS as models of established infectious disease. More recently, cryopreserved uninfected PCLS have been thawed and subsequently infected with influenza virus to investigate virus-host interactions (36). In contrast, the present study demonstrates that PCLS generated from mice with established *P. murina* infection prior to cryopreservation maintain both host tissue viability and sufficient *P. murina* viability to support antifungal efficacy testing while preserving the native infected lung microenvironment.

Our findings expand the utility of cryopreserved PCLS beyond studies of pulmonary physiology and toxicology to investigations of fungal host-pathogen interactions and preclinical antifungal drug evaluation. Specifically, we show that cryopreserved PCP-infected PCLS maintain tissue integrity, host cell viability, and sufficient *P. murina* viability to support molecular analyses and antifungal efficacy studies following thawing. Collectively, these findings establish cryopreserved PCP-infected PCLS as a practical, reproducible, and biologically relevant ex vivo platform for investigating *Pneumocystis* biology and evaluating candidate antifungal therapies.

As expected, cryopreservation resulted in some reduction in overall metabolic activity, as measured by the resazurin reduction assay. Nevertheless, both uninfected and infected-PCP thawed PCLS retained reasonable levels of measurable metabolic activity, indicating that the lung slices remained viable following cryopreservation. Similar reductions in metabolic activity have been reported following cryopreservation of PCLS from multiple species and are generally attributed to cellular stress associated with freezing, ice crystal formation, and osmotic injury during thawing rather than complete tissue loss (34, 37–39). Importantly, although cryopreservation somewhat reduced metabolic activity, it did not abolish tissue or fungal viability, supporting continued use of the PCP-infected slices for downstream experimental applications.

The preservation of infection-associated epithelial gene expression provided additional evidence that cryopreservation maintains the biological characteristics of PCP-infected PCLS. Consistent with previous reports demonstrating that *Pneumocystis* infection disrupts alveolar epithelial homeostasis and surfactant biology (40–43), PCP-infected PCLS exhibited increased *Pdpn* expression and reduced *Sftpc* expression compared with uninfected controls. These infection-associated expression patterns were maintained following cryopreservation, despite an overall reduction in tissue metabolic activity, indicating that freezing and thawing did not substantially alter the epithelial response to *P. murina* infection. Although *Pdpn* and *Sftpc* are established markers of alveolar type I and type II epithelial cells, respectively (44–47), the present study was not designed to define the cellular mechanisms underlying these transcriptional changes. Rather, preservation of these epithelial gene expression patterns, together with maintained tissue viability, ultrastructural integrity, and responsiveness to pentamidine, further supports the utility of cryopreserved PCP-infected PCLS as a physiologically relevant ex vivo platform for studying *Pneumocystis* biology and evaluating antifungal therapies.

Following characterization of tissue viability and epithelial gene expression, we sought to determine whether cryopreserved PCP-infected PCLS remained suitable for antifungal efficacy testing. Although cryopreservation significantly reduced overall metabolic activity and *P. murina* viability compared with freshly prepared infected PCLS, viable organisms remained detectable following thawing. Importantly, treatment of post-thaw PCP-infected PCLS with pentamidine for 72 h significantly reduced *P. murina* viability compared with DMSO-treated controls, demonstrating that cryopreserved infected PCLS retain sufficient viable organisms to support antifungal efficacy studies. Pentamidine is a well-established anti-*Pneumocystis* agent that has demonstrated efficacy in both experimental models and the clinical treatment of PCP (27, 48–50), making it an appropriate positive-control compound for proof-of-concept evaluation of the cryopreserved PCLS model. The reduction in *P. murina* viability observed in our study is therefore consistent with its established anti-*Pneumocystis* activity and provides biological validation of the cryopreserved PCLS model. Importantly, pentamidine treatment did not significantly alter resazurin reduction in either uninfected or PCP-infected PCLS, indicating that the concentration used did not measurably impair host tissue metabolic activity during the 72-h treatment period. This finding suggests that the reduction in *P. murina* viability reflects the expected anti-*Pneumocystis* activity of pentamidine rather than nonspecific toxicity to the lung slices. Together, these findings demonstrate that cryopreserved PCP-infected PCLS retain both host tissue integrity and sufficient pathogen viability to support pharmacological evaluation of antifungal compounds following thawing.

The ability to cryopreserve infected PCLS offers several practical advantages over freshly prepared tissue. Cryopreservation enables infected lung slices to be generated in advance, stored long term, and thawed as needed, thereby reducing dependence on continuous animal infections and minimizing experimental variability associated with infection burden between animals (51). In addition, this approach facilitates standardized drug efficacy studies using biologically comparable infected tissue across multiple experiments, investigators, and laboratories. To our knowledge, this is the first study demonstrating that PCLS generated from mice with established PCP prior to cryopreservation remain suitable for molecular analyses and antifungal efficacy testing following thawing. Whereas previous studies have used cryopreserved uninfected PCLS for pulmonary physiology, toxicology, and experimental pathogen challenge, our findings extend the application of cryopreserved PCLS to the study of established fungal infection and preclinical antifungal drug evaluation. These advantages have the potential to facilitate multicenter studies and higher-throughput evaluation of candidate antifungal compounds while reducing animal use.

Several limitations should be acknowledged. This study evaluated a single cryopreservation time interval and a single antifungal agent. Pentamidine was selected for these proof-of-concept studies because of its well-established anti-*Pneumocystis* activity in vitro and its suitability as a positive-control agent for evaluating antifungal efficacy in the cryopreserved PCLS model (27, 52, 53). Future studies will evaluate trimethoprim-sulfamethoxazole (TMP-SMX), the current standard-of-care therapy for PCP (54), to further validate the utility of cryopreserved PCLS for antifungal efficacy testing. Because TMP-SMX exerts its antimicrobial activity by sequential inhibition of folate biosynthesis and is typically administered over longer treatment durations in both experimental models (55) and clinical practice (56), pentamidine provided a more practical agent for the initial in vitro validation of this cryopreserved PCLS platform (50, 57). Additional studies examining longer cryopreservation intervals, alternative cryoprotective formulations, and other clinically relevant antifungal compounds will further define the utility of this model. Likewise, although epithelial gene expression and tissue morphology were preserved, comprehensive transcriptomic and functional analyses (58) will be valuable for determining the extent to which cryopreserved PCLS recapitulate fresh tissue responses over prolonged culture periods.

In conclusion, we demonstrate that PCP-infected PCLS remain structurally intact, metabolically active, and transcriptionally responsive following cryopreservation while retaining sufficient *P. murina* viability for antifungal efficacy testing. These findings establish cryopreserved PCP-infected PCLS as a robust and practical ex vivo platform for studying *Pneumocystis* host-pathogen interactions and evaluating novel therapeutic strategies.

## ETHICAL STATEMENT

This study was performed in strict accordance with the recommendations in the Guide for the Care and Use of Laboratory Animals (59). The Mayo Clinic Institutional Animal Care and Use Committee approved this research (A00008661-26).

## Supplementary Material

**Supplementary Table S1.**
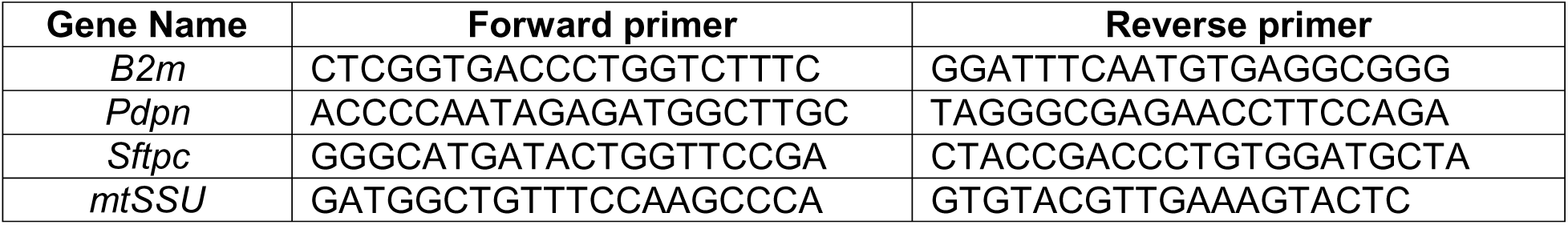
qPCR primer list.

